# Respiration, heartbeat, and conscious tactile perception

**DOI:** 10.1101/2021.03.22.436396

**Authors:** Martin Grund, Esra Al, Marc Pabst, Alice Dabbagh, Tilman Stephani, Till Nierhaus, Michael Gaebler, Arno Villringer

**Author notes:** Corresponding author: Martin Grund, Max Planck Institute for Human Cognitive and Brain Sciences, Stephanstr. 1A, 04103 Leipzig, Germany. **Conflict of Interest:** The authors declare no competing financial interest.

## Abstract

Previous studies have shown that timing of sensory stimulation during the cardiac cycle interacts with perception. Given the natural coupling of respiration and cardiac activity, we investigated here their joint effects on tactile perception. Forty-one healthy female and male human participants reported conscious perception of finger near-threshold electrical pulses (33% null trials) and decision confidence while electrocardiography, respiratory activity, and finger photoplethysmography were recorded. Participants adapted their respiratory cycle to expected stimulus onsets to preferentially occur during late inspiration / early expiration. This closely matched heart rate variation (sinus arrhythmia) across the respiratory cycle such that most frequent stimulation onsets occurred during the period of highest heart rate probably indicating highest alertness and cortical excitability. Tactile detection rate was highest during the first quadrant after expiration onset. Inter-individually, stronger respiratory phase-locking to the task was associated with higher detection rates. Regarding the cardiac cycle, we confirmed previous findings that tactile detection rate was higher during diastole than systole and newly specified its minimum at 250 - 300 ms after the R-peak corresponding to the pulse wave arrival in the finger. Expectation of stimulation induced a transient heart deceleration which was more pronounced for unconfident decision ratings. Inter-individually, stronger post-stimulus modulations of heart rate were linked to higher detection rates. In summary, we demonstrate how tuning to the respiratory cycle and integration of respiratory-cardiac signals are used to optimize performance of a tactile detection task.

**Significance statement:** Mechanistic studies on perception and cognition tend to focus on the brain neglecting contributions of the body. Here, we investigated how respiration and heartbeat influence tactile perception: Respiration phase-locking to expected stimulus onsets corresponds to highest heart rate (and presumably alertness/cortical excitability) and correlates with detection performance. Tactile detection varies across the heart cycle with a minimum when the pulse reaches the finger and a maximum in diastole. Taken together with our previous finding of unchanged early ERPs across the cardiac cycle we conclude that these effects are not a peripheral physiological artifact but a result of cognitive processes that model our body’s internal state, make predictions to guide behavior, and might also tune respiration to serve the task.

## Introduction

Our body senses signals from the outer world (exteroception), but also visceral signals from inside the body (interoception) and it has been shown that these two continuous types of perception interact (Critchley and Harrison, 2013; Critchley and Garfinkel, 2015; Babo-Rebelo et al., 2016; Seth and Friston, 2016; Azzalini et al., 2019). For example, we have recently shown that tactile perception interacts with cardiac activity as conscious detection of near-threshold stimuli was more likely towards the end of the cardiac cycle (Motyka et al., 2019; Al et al., 2020) and was followed by a more pronounced deceleration of heart rate as compared to missed stimuli (Motyka et al., 2019). In line with increased detection during diastole, late (P300) cortical somatosensory evoked potentials (SEPs) were also higher during diastole as compared to systole (Al et al., 2020). A similar cardiac phase-dependency has also been revealed for visual sampling: microsaccades and saccades were more likely during systole, whereas fixations and blinks during diastole (Ohl et al., 2016; Galvez-Pol et al., 2020). Following an interoceptive predictive coding account, the very same brain model that predicts (e.g., cardiac-associated) bodily changes and suppresses their access to consciousness (as unwanted “noise”) might also suppress perception of external stimuli which happen to coincide with those bodily changes (Allen et al., 2019).

Another dominant body rhythm that can even be regulated intentionally in contrast to cardiac activity is the respiration rhythm (Azzalini et al., 2019). Also for respiration, which naturally drives and is driven by cardiac activity (Kralemann et al., 2013; Dick et al., 2014), phase-dependency of behavior and perception has been reported. For instance, self-initiated actions were more likely during expiration, whereas externally-triggered actions showed no correlation with the respiration phase (Park et al., 2020). Furthermore, inspiration onsets were reported to be phase-locked to task onsets which resulted in greater task-related brain activity and increased task performance for visuospatial perception, memory encoding and retrieval, and fearful face detection (Huijbers et al., 2014; Zelano et al., 2016; Perl et al., 2019). Respiratory phase locking has also been shown for brain rhythms and cortical excitability offering a neurophysiological basis for modulations of task performance and accordingly, phase locking to respiration has been interpreted as tuning the sensory system for upcoming information (Perl et al., 2019). Thus, for both, the cardiac cycle and the respiratory cycle, certain phases might also be beneficial for conscious perception and could be timed to paradigms instead of modelled as noise within an interoceptive predictive coding framework. While cardiac activity and respiration are closely interdependent, it remains unclear how they jointly shape perceptual processes.

Our present study combined the observation of cardiac and respiratory activity with a paradigm that asked participants to report (a) detection of weak electrical pulses applied to their left index finger and (b) their decision confidence. Decision confidence was assessed to identify the link between metacognition and cardiorespiratory cycle effects on somatosensation. As we have previously shown that greater tactile detection during diastole corresponded to increased perceptual sensitivity and not to a more liberal response criterion (Al et al., 2020), we expected the cardiac cycle effect not to be a side-effect of unconfident perceptual decisions. Afferent fibers in the finger have been reported to be modulated by cardiac pressure changes which the brain has to ignore or filter out (Macefield, 2003). Thus, we measured photoplethysmography to investigate whether cardiac related movements in the finger caused by the blood pulse wave coincided with lower tactile detection during systole. Furthermore, we tried to capture early SEPs at the upper arm to rule out differences in (peripheral) SEP amplitudes as explanation for altered conscious tactile perception across cardiac or respiratory cycles.

This study setup was intended to address the following research questions:

- Does the interaction of cardiac activity and conscious tactile perception depend on decision confidence?
- What is the precise temporal relationship between suppressed tactile detection and the kinetics of the pulse wave in the finger?
- Does conscious tactile perception vary across the respiratory cycle and what is the relationship to respiratory modulation of the heartbeat?

## Methods

### Participants

Forty-one healthy humans (21 women, mean age = 25.5, age range: 19-37) participated in the study. Participants were predominantly right-handed with a mean laterality index of 90, *SD* = 17 (Oldfield, 1971). For four participants, the mean laterality index was not available.

### Ethics statement

All participants provided an informed consent. The experimental procedure and physiological measurements were approved by the ethics commission at the medical faculty of the University of Leipzig.

### Experimental design and statistical analysis

The experiment was designed to capture tactile detection of near-threshold stimuli (50% detection) and trials without stimuli (0% detection) across the cardiac and respiratory cycle. This resulted in three main stimulus-response conditions: (a) correct rejections of trials without stimulation, (b) undetected (misses), and (c) detected near-threshold stimuli (hits). False alarms (yes-responses) during trials without stimulation were very rare (mean FAR = 6%, *SD* = 6%) and thus not further analyzed. Additionally, participants reported their decision confidence which allowed us to split trials by confidence.

We applied circular statistics to investigate whether conscious tactile perception was uniformly distributed across the cardiac and respiratory cycle or showed unimodal patterns. For each stimulus onset, the temporal distances to the preceding R-peak and expiration onset were calculated and put in relation to its current cardiac and respiratory cycle duration measured in degrees. Following for each participant, these angles were averaged for hits, misses, and correct rejections. For each stimulus-response condition, the resulting distributions of mean angles were tested across participants with the Rayleigh test for uniformity (Landler et al., 2018) from the R package “circular” (Version 0.4-93). The application of circular statistics had two advantages: First, it accounted for cardiac and respiratory cycle duration variance within and between participants. Second, it allowed us to determine phases when detection differed without having to rely on arbitrary binning. However, it assumed that the different phases of the cardiac and respiratory cycle behave proportionally the same when cycle duration changes. That is why we complemented the circular statistics with a binning analysis that investigated the near-threshold detection rate for fixed time intervals relative to the preceding R-peak and expiration onset.

In repeated-measures ANOVAs, Greenhouse-Geisser correction was used to adjust for the lack of sphericity. Post-hoc *t*-tests p-values were corrected for multiple comparisons with a false discovery rate (FDR) of 5% (Benjamini and Hochberg, 1995).

### Data and code availability

The code to run and analyze the experiment is available at http://github.com/grundm/respirationCA. The behavioral and physiological data

(electrocardiogram, respiration, and oximetry) can be shared by the corresponding author upon request if data privacy can be guaranteed.

### Stimuli and apparatus

Somatosensory stimulation was delivered via steel wire ring electrodes to the left index finger with a constant current stimulator (DS5; Digitimer, United Kingdom). The anode was placed on the middle phalanx and the cathode on the proximal phalanx. The stimuli were single square-wave pulses with a duration of 0.2 ms and a near-threshold intensity (50% detection rate) which was assessed prior to an experimental block with an automatic procedure as described in the last paragraph of the section Behavioral paradigm. The stimulator was controlled by the waveform generator NI USB-6343 (National Instruments, Austin, Texas) and custom MATLAB scripts using the Data Acquisition Toolbox (The MathWorks Inc., Natick, Massachusetts, USA).

### Behavioral paradigm

Participants had to report whether they perceived an electrical pulse and whether this yes/no-decision was confident or not. The experiment was separated into four blocks. Each block consisted of 150 trials. Participants received a near-threshold stimulus in 100 trials (mean intensity = 1.96 mA, range: 0.76-3.22 mA). In 50 trials, there was no stimulation (33% catch trials). The order of near-threshold and catch trials was pseudo-randomized for each block and participant. In total, there were 400 near-threshold and 200 catch trials.

Each trial started with a black fixation cross (black “+”) for a counterbalanced duration of 1.0-2.0 s (Figure 1). It was followed by a salmon-colored fixation cross (0.62 s) to cue the stimulation at 0.5 s after the cue onset. With the cue offset, the participants had to report the detection of a tactile stimulus (yes/no). After the yes/no-button press, a pause screen was displayed for 0.3 s, before the participants were asked to report their decision confidence (confident/unconfident). With pressing the button for “confident” or “unconfident”, the new trial started. For both reports, the maximum response time was 1.5 s. Thus, the maximum possible trial duration was 5.8 s.

**Figure 1.**
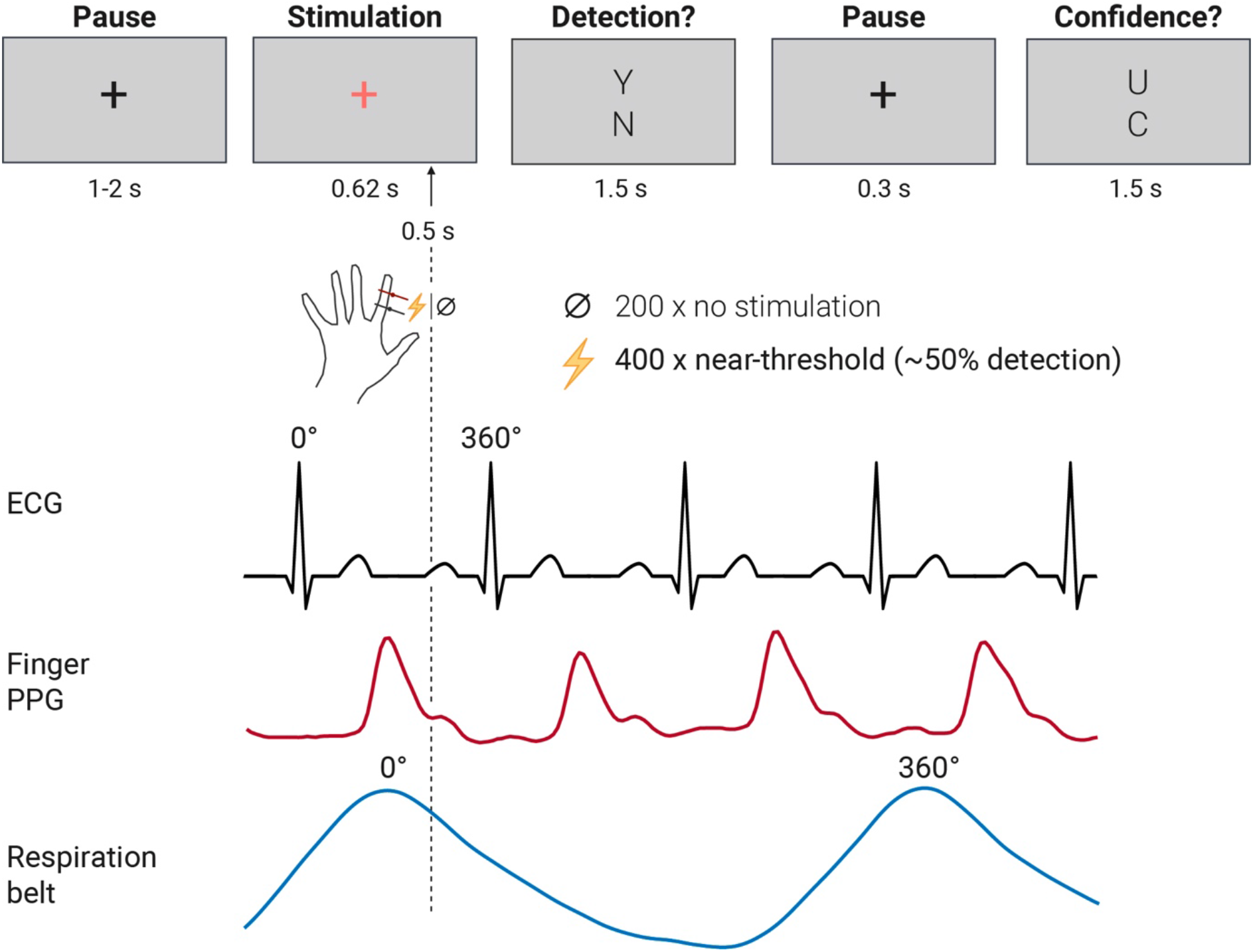
Experimental procedure and physiological parameters visualized for one exemplary trial. The tiles represent the participant’s visual display and the times given below indicate the presentation duration. The near-threshold electrical finger nerve stimulation was always 0.5 s after the cue onset (salmon-colored fixation cross). Here only one of four button response mappings is displayed (Y = yes; N = no; U = unconfident; C = confident). In total, 400 near-threshold trials and 200 trials without stimulation (33% catch trials) were presented in randomized order. Exemplary traces of electrocardiogram (ECG), finger photoplethysmography (PPG), and respiration belt below the trial procedure indicate that stimulus detection was analyzed relative to cardiac and respiratory cycles (0-360°).

Participants indicated their perception and decision confidence with the right index finger on a two-button box. The buttons were arranged vertically. The four possible button response mappings were counterbalanced across participants, so that the top button could be assigned to “yes” or “no”, and “confident” or “unconfident” respectively for one participant.

Prior to the experiment, participants were familiarized with the electrical finger nerve stimulation and an automatic threshold assessment was performed in order to determine the stimulus intensity corresponding to the sensory threshold (50% detection rate). The threshold assessment entailed an up-and-down procedure (40 trials in the first run and 25 trials in subsequent runs) which served as a prior for the following Bayesian approach (psi method; 45 trials in first run and 25 trials in subsequent runs) from the Palamedes Toolbox (Kingdom and Prins, 2009) and closed with a test block (5 trials without stimulation and 10 trials with stimulation intensity at the threshold estimate by psi method). Based on the test block results for the psi method threshold estimate and weighting in the results of the up-and-down procedure, the experimenter selected a near-threshold intensity candidate for the first experimental block. The visual display of the trials in the threshold assessment was similar to the trials in the experimental block (Figure 1) but without the confidence rating and a shorter fixed intertrial interval (0.5 s). If a block resulted in a detection rate diverging strongly from 50% (smaller than 25% or greater than 75%), the threshold assessment was repeated before the subsequent block to ensure a detection rate of about 50% throughout the whole experiment. The experimental procedure was controlled by custom MATLAB scripts using the Psychophysics Toolbox (Kleiner et al., 2007).

### Electrocardiogram acquisition

The electrocardiogram (ECG) was recorded with the BrainAmp ExG (Brain Products, Gilching, Germany) between two adhesive electrodes that were placed on the sternum and just below the heart on the left side of the thorax. A ground electrode was placed on the acromion of the right shoulder. The sampling frequency was 5000 Hz for 39 participants. Two participants were recorded with 1000 Hz.

### Respiration acquisition

Respiration was measured with a respiration belt (BrainAmp ExG; Brain Products, Gilching, Germany). The belt with a pressure-sensitive cushion was placed at the largest expansion of the abdomen during inspiration. The sampling frequency was 5000 Hz for 39 participants. Two participants were recorded with 1000 Hz.

### Peripheral nerve activity acquisition

To examine the possibility to measure somatosensory evoked potentials (SEP) of peripheral nerve activity in response to near-threshold finger stimulation, two surface electrodes were placed with a distance of 2 cm at the left upper inner arm (below the biceps brachii) above the pathway of the median nerve in a sub-sample of 12 participants. The signal was recorded with a sampling rate of 5000 Hz, low-pass filtered at 1000 Hz, using a bipolar electrode montage (BrainAmp ExG; Brain Products, Gilching, Germany).

### Oximetry acquisition

The photoplethysmography was recorded with a finger clip SpO_2_ transducer at the left middle finger at 50 Hz (OXI100C and MP150; BIOPAC Systems Inc., Goleta, California, USA).

### Behavioral data analysis

The behavioral data was analyzed with R 4.0.3 in RStudio 1.3.10923. First, trials were filtered for detection and confidence responses within the maximum response time of 1.5 s. Second, only blocks were considered with a near-threshold hit rate at least five percentage points above the false alarm rate. These resulted in 37 participants with 4 valid blocks, 2 participants with 3 valid blocks and 2 participants with 2 valid blocks. The frequencies of the response “confident” for correct rejections, misses, and hits were compared with paired *t*-tests. Furthermore, the detection and confidence response times and resulting trial durations were compared between correct rejections, misses, and hits with paired *t*-tests. The response times for hits and miss were additionally compared between confident and unconfident near-threshold trials.

### Cardiac data analysis

ECG data was preprocessed with Kubios (Version 2.2) to detect R-peaks. For two participants, the first four and the first twenty-two trials respectively had to be excluded due to a delayed start of the ECG recording. Additionally, one block of one participant and two blocks of another participant were excluded due to corrupted ECG data quality based on visual inspection (no R-peak detection possible).

First for correct rejections, misses, and hits, the circular distribution within the cardiac cycle was assessed with the Rayleigh test of uniformity and compared between the stimulus-response conditions with a randomization version of Moore’s test for paired circular data (Moore, 1980) based on 10,000 permutations as implemented by (Pewsey et al., 2013). Additionally, this analysis was repeated for confident and unconfident hits and misses.

Second, instead of the relative position within the cardiac cycle, near-threshold trials were binned to four time intervals based on their temporal distance from the previous R-peak (0 - 200 ms, 200 - 400 ms, 400 - 600 ms, and 600 - 800 ms). Then, dependent probabilities were calculated for each of the four possible outcomes (unconfident misses, confident misses, unconfident hits, and confident hits) given the time interval. The probabilities were compared with *t*-tests between time intervals separately for each of the four possible outcomes. FDR-correction was applied across all 24 *t*-tests. Furthermore, we used linear mixed-effects models (LMMs) with maximum likelihood estimation (Laplace approximation; glmer function in R) to evaluate the cardiac cycle effect on confidence beyond detection in near-threshold trials. Models without (“confidence ∼ detection + (1|participant)”) and with the cardiac cycle information (“confidence ∼ detection + cos(cardiac_phase) + sin(cardiac_phase) + (1|participant)”) were then compared using the anova function in R.

Additionally, we assessed whether metacognition changed across the cardiac cycle. For this purpose, response-specific meta-*d’* was estimated for each cardiac interval using a hierarchical Bayesian model (MATLAB function fit_meta_d_mcmc_group.m) from the HMeta-d toolbox (Maniscalco and Lau, 2012; Fleming, 2017). Complementary to *d’* for yes/no-detection tasks, meta-*d’* measures the sensitivity to distinguish with confidence ratings correct from incorrect yes/no-decisions and is calculated separately for yes and no responses (false alarms vs. hits; misses vs. correct rejections). For estimating meta-*d’*, the Markov chain Monte Carlo (MCMC) method used three chains of 10.000 iterations with 1000 burn-in samples and default initial values from JAGS. We also ensured that 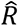 was < 1.1 for all model parameters of interest, indicating convergence of the model fit (Fleming, 2017). To account for perceptual sensitivity *d’* in measuring metacognitive sensitivity meta-*d’*, M-ratios were calculated (meta-*d’* divided by *d’* given yes or no response). An M-ratio of 1 indicates that confidence ratings can perfectly discriminate between correct and incorrect responses (Fleming and Lau, 2014). Compared to model-free approaches, M-ratio controls for differences in first order performance (*d’*) as well as response biases (*c*). For statistical testing whether M-ratios differed between yes/no-responses or cardiac cycle intervals, the differences of the corresponding group-level M-ratio posterior distributions were calculated and assessed whether their 95% high-density intervals entailed zero or not (Kruschke, 2015; Fleming, 2017). The latter was interpreted as evidence for a difference.

Third, we analyzed the interbeat intervals in the course of a trial between the stimulus-response conditions. For this, two interbeat intervals before, one during, and two after the stimulus onset were selected and compared with a two-way repeated measures ANOVA and post-hoc *t*-tests. To investigate whether changes of the interbeat interval length were caused by systole or diastole, we applied a trapezoidal area algorithm (Vázquez-Seisdedos et al., 2011) to detect T-waves within the first post-stimulus interbeat interval (“S+1”). The T-wave end defines the end of systole and the onset of diastole. Systole and diastole durations were averaged for each participant and stimulus-response condition (correct rejection, miss, hit) and compared between the conditions across participants with FDR-corrected *t*-tests. After visual inspection of T-wave detection results, two participants were excluded from the analysis, because the cardiac signal did not allow to consistently detect the T-wave. Additionally, in two trials of one participant the t-wave detection was not successful and 127 trials with a systole length three standard deviations below or above the participant mean at “S+1” were excluded.

Furthermore, the relationship between heart slowing and detection performance was analyzed by calculating Pearson’s correlation coefficients between heart slowing (ratio from interbeat interval “Stimulus” to “S+1”) and near-threshold detection rate for (a) all trials and (b) correct rejections.

### Oximetry data analysis

Oximetry data was analyzed with custom MATLAB scripts to detect pulse wave peaks with a minimum peak distance based on 140 heartbeats per minute (25.7 s) and a minimum peak prominence equal to a tenth of the data range in each block. Pulse wave cycles with a duration 1.5 times the median duration of the respective block were excluded from further processing. In R, pulse wave cycle data were merged with the behavioral data to apply the same exclusion criteria. Pulse wave peaks were located in the cardiac cycle to assess the duration since the previous R-peak (pulse wave transit time, PWTT) and its relative position in degree within the cardiac cycle.

### Respiration data analysis

After visual inspection of the respiration traces, respiratory cycle detection was performed following the procedure by Power et al. (2020). First, outliers were replaced in a moving 1-s window with linearly interpolated values based on neighboring, non-outlier values. Local outliers were defined as values of more than three local scaled median absolute deviations (MAD) away from the local median within a 1-s window (Power et al., 2020). MAD was chosen for its robustness compared to standard deviation which is more affected by extreme values. Subsequently, the data was smoothed with a 1-s window Savitzky-Golay filter (Savitzky and Golay, 1964) to facilitate peak detection. Traces were then z-scored to identify local maxima (inspiration onsets) and in the inverted trace local minima (expiration onsets) with the MATLAB findpeaks function. Local maxima and minima had to be at least 2 s apart with a minimum prominence of 0.9 times the interquartile range of the z-scored data. Respiratory cycles were defined as the interval from one expiration onset to the next expiration onset. For each participant, respiratory cycles with more than two times the median cycle duration were excluded from further analysis.

For each stimulus-response condition and participant, the mean angle direction of stimulus onsets within the respiratory cycle and their circular variance across trials were calculated. The distribution of mean angles of each stimulus-response condition was tested for uniformity with the Rayleigh test. Circular variance was defined as *V* = 1-*R*, where *R* is the mean resultant length of each stimulus-response condition and participant with values between 0 and 1. Differences in circular variances between stimulus-response conditions were assessed with paired *t*-tests. To investigate whether respiration phase-locking showed a relationship with task performance, Pearson’s correlation coefficients between circular variance of respiration phases and near-threshold detection rate was calculated in (a) in all trials and (b) correct rejections.

Furthermore, we investigated whether participants gradually aligned their respiration to the stimulus onset in the beginning of the experiment. For the first 30 trials, the difference between each trial’s stimulus onset angle and the mean angle within the first block was determined (“diff_angle2mean”). The trial angle difference from the mean was used as a dependent variable in a random-intercept linear regression based on maximum likelihood estimation with trial number as independent variable: “diff_angle2mean ∼ 1 + trial + (1|participant)”. The fit of this model was compared with a random-intercept only model “diff_angle2mean ∼ 1 + (1|participant)” in a *χ*²-test to assess the effect of trial number on the angle difference. This analysis included only the 37 participants with a valid first block and excluded trials with false alarms.

Heart rate was analyzed across the respiratory cycle by assigning trials according to their respiration phase at stimulus onset to eight 45°-intervals. For each interval the corresponding cardiac interbeat intervals at stimulus onset were averaged for each participant and compared with FDR-corrected *t*-tests.

Lastly, we compared the respiratory cycle duration between stimulus-response conditions by performing a one-way repeated-measures ANOVA and post-hoc *t*-tests. Furthermore, the inspiration onset within each respiratory cycle was determined to statistically compare expiration and inspiration duration between stimulus-response conditions with a two-way repeated measures ANOVA.

### Phase-locking analysis between cardiac and respiratory activity

The n:m (*n*, *m* ∈ *N*) synchronization (Lachaux et al., 1999) was calculated in an inter-trial setting for the stimulation onset as the following:

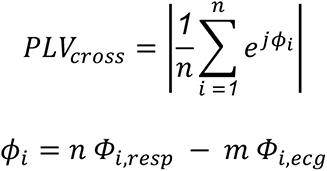

where *Φ_i,resp_* and *Φ_i,ecg_* were the stimulation onset angles for the i-th trial within the respiratory (*resp*) and cardiac cycle (*ecg*), and *j* was the imaginary number. While *m* = *1* was chosen for all participants, values for *n* were selected by calculating the ratio of the cardiac and respiratory frequency rounded to the nearest integer. The frequencies were estimated based on the mean cardiac and respiratory cycle durations at stimulus onset. The inter-trial n:m synchronization at stimulation onset can provide information about the extent to which the weighted phase difference of the two signals stays identical over trials. The calculated phase-locking value (PLV) lies between zero and one, with zero indicating no inter-trial coupling and one showing a constant weighted phase-difference of the two signals at the stimulation time.

### Somatosensory evoked potential analysis

For the twelve participants with peripheral nerve recordings, stimulation artefacts were removed with a cubic monotonous Hermite spline interpolation from -2 s until 4 s relative to the trigger. Next, a 70-Hz high-pass filter was applied (4th order Butterworth filter applied forwards and backwards) and the data was epoched - 100 ms to 100 ms relative to the trigger with -50 ms to -2 ms as baseline correction. Subsequently, epochs were averaged across for valid trials (yes/no and confidence response within maximum response time) with near-threshold stimuli and without stimulations.

## Results

### Detection and confidence responses

Participants (*N* = 41) detected on average 51% of near-threshold stimuli (*SD* = 16%) and correctly rejected 94% of catch trials without stimulation (*SD* = 6%). On average, 188 catch trials (range: 93-200) and 375 near-threshold trials (range: 191-400) were observed. Participants reported to be “confident” about their yes/no-decision in 88% of the correct rejections (*SD* = 13%), in 71% of the misses (*SD* = 21%), and in 62% of the hits (*SD* = 18%). The confidence rate differed significantly between all conditions in paired *t*-tests (CR vs. miss: *p* = 3 * 10^-8^; CR vs. hit: *p* = 1 * 10^-9^; miss vs. hit: *p* = 0.019). In total, we observed on average 184 misses (range: 58-303), 192 hits (range: 59-302), 177 correct rejections (range: 72-198) and 11 false alarms (range: 0-36). Two-third of the participants (27) had less than ten false alarms and four participants had zero false alarms. Due to zero or very few observations, false alarms were not further analyzed.

In near-threshold trials, participants reported their yes/no-decision later than for correct rejections (mean ± SD: *RT_Hit_* = 641±12 ms, *RT_Miss_* = 647±12 ms, *RT_CR_* = 594±10 ms; paired *t*-test hit vs. CR: *p* = 0.02, miss vs. CR: *p* = 3 * 10^-8^). The yes/no-response times for hits and misses did not differ significantly (*p* = 0.43). Additionally in unconfident compared to confident near-threshold trials, yes/no-responses were on average 221 ms slower (mean ± SD: *RT_Near_unconf_* = 789±11 ms, *RT_Near_conf_* = 569±9 ms; paired *t*-test: *p* = 2 * 10^-16^). Splitting near-threshold trials by confidence resulted in on average 49 unconfident misses (range: 6-143), 135 confident misses (range: 29-289), 70 unconfident hits (range: 9-181), and 122 confident hits (range: 24-277).

### Cardiac cycle

First, we addressed the question whether stimulus detection differed along the cardiac cycle. For hits, mean angles within the cardiac cycle were not uniformly distributed (*R* = 0.34, *p* = 0.007; Figure 2), indicating a relation between cardiac phase and stimulus detection. Sixteen participants had a mean angle for hits in the last quarter of the cardiac cycle (270-360°). The Rayleigh tests were not significant for misses (*R* = 0.20, *p* = 0.18) and correct rejections (*R* = 0.05, *p* = 0.91). Additionally, we tested whether there was a bias for the presentation of near-threshold stimuli within the cardiac cycle and calculated a Rayleigh test for each participant. None of these tests was significant (all FDR-corrected *p* > 0.31). With a randomization version of Moore’s test for paired circular data based on 10,000 permutations the distributions of correct rejections, misses, and hits were statistically compared to each other. The distribution of hits and misses differed significantly (*R* = 1.27, *p* = 0.01), whereas the distributions of correct rejections and misses (*R* = 0.84, p = 0.13), and correct rejections and hits did not (*R* = 0.41, *p* = 0.62).

**Figure 2.**
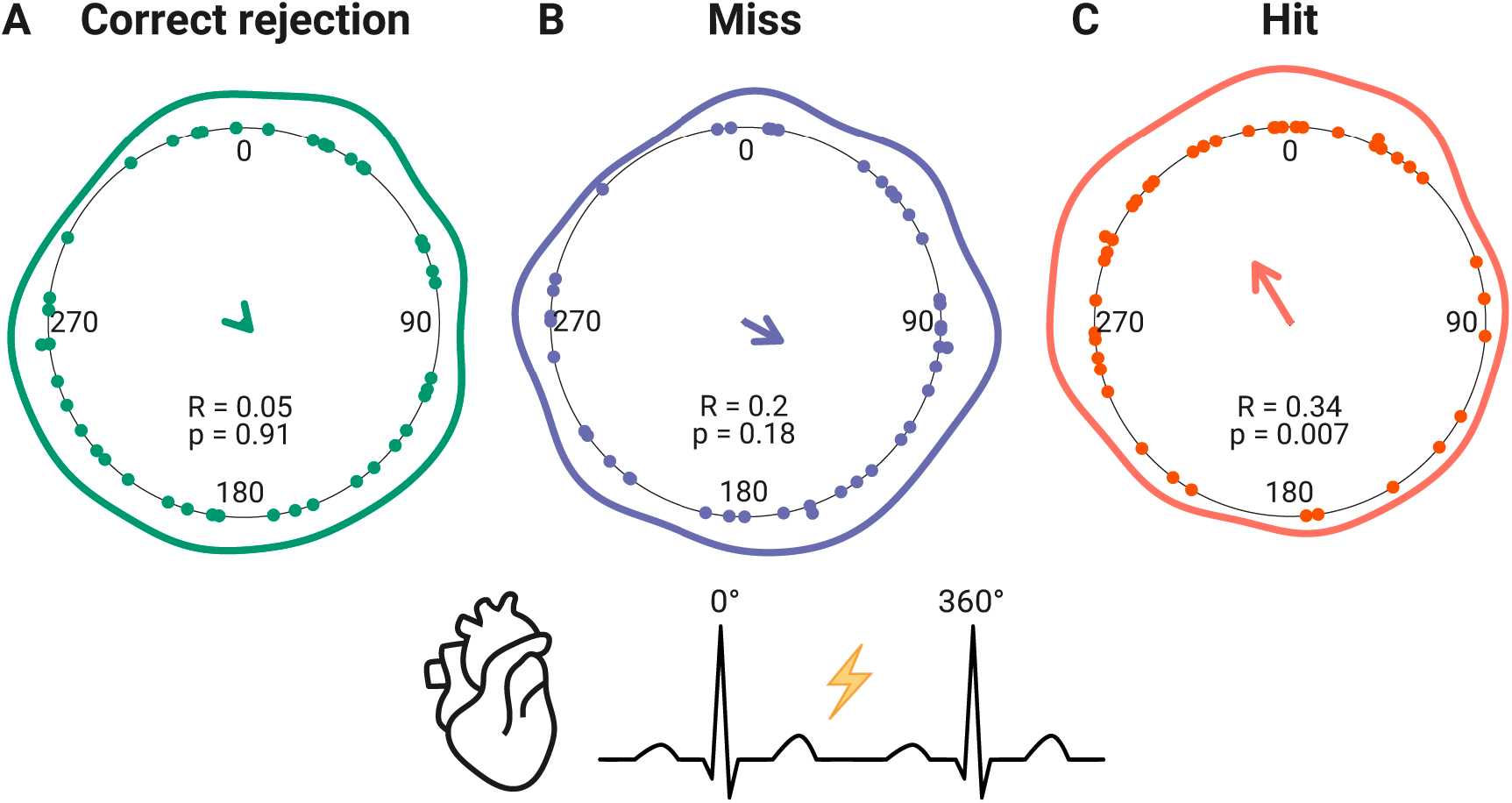
Distribution of mean angles (stimulus onset relative to cardiac cycle) for (***A***) correct rejections (green), (***B***) misses (purple), and (***C***) hits (red). Each dot indicates the mean angle of one participant. The line around the inner circle shows the density distribution of these mean angles. The direction of the arrow in the center indicates the mean angle across the participants while the arrow length represents the mean resultant length *R*. The resulting p-value of the Rayleigh test of uniformity is noted below.

**Figure 3.**
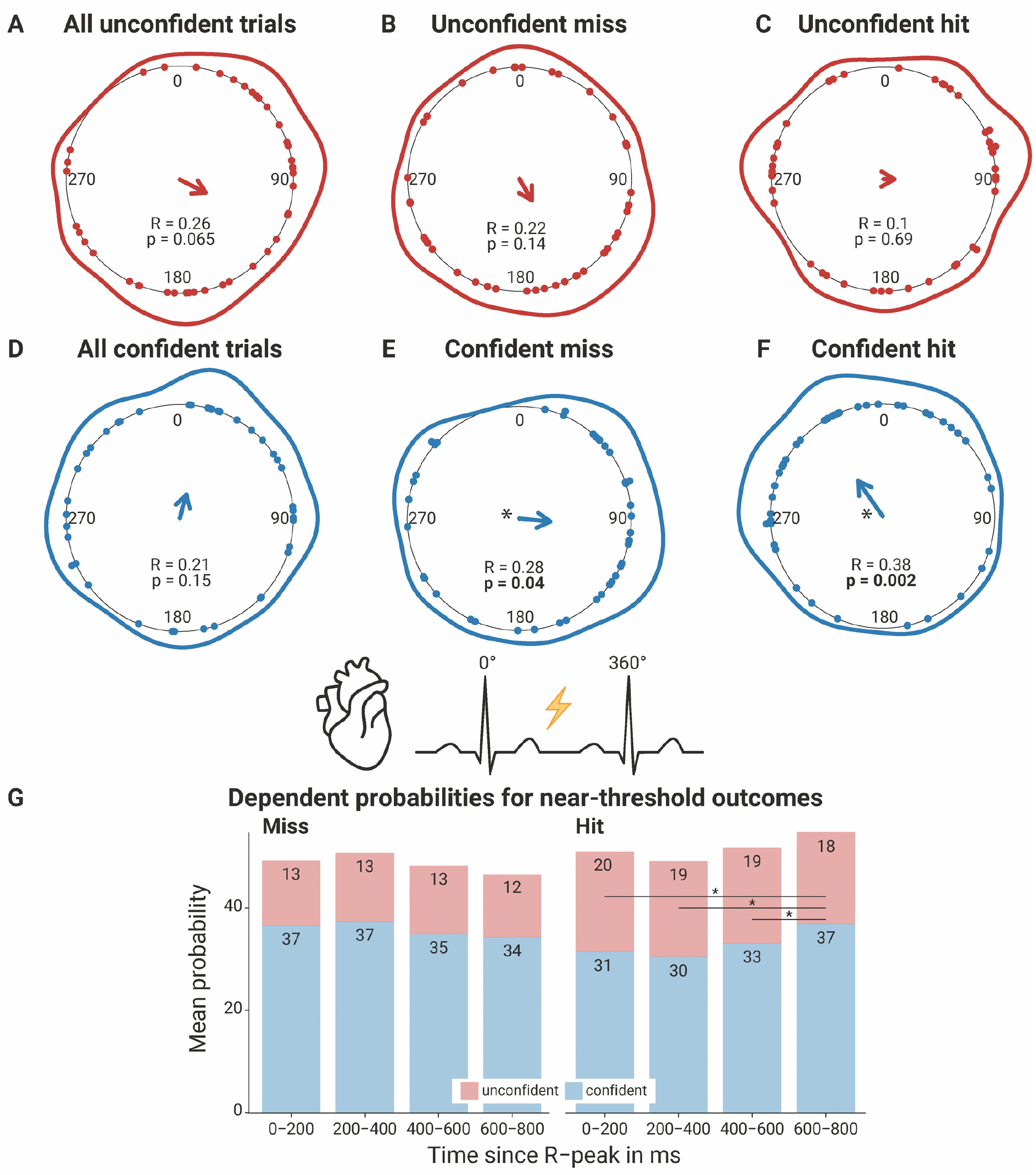
Circular distribution within the cardiac cycle of unconfident/confident trials and unconfident/confident misses and hits (***A-F***), dependent probabilities of unconfident/confident miss/hit at four time intervals after the R-peak (***G***), and metacognitive efficiency across the cardiac cycle (***H***). The distributions of mean angles (stimulus onset relative to cardiac cycle) are shown for (***A***) all unconfident trials (correct rejections, misses, and hits), (***B***) unconfident misses (red), (***C***) unconfident hits (red), (***D***) all confident trials (correct rejections, misses, and hits), (***E***) confident misses (blue), and (***F***) confident hits (blue). In ***A-F***, each dot indicates the mean angle of one participant. The line around the inner circle shows the density distribution of these mean angles. The direction of the arrow in the center indicates the mean angle across the participants while the arrow length represents the mean resultant length *R*. The resulting p-value of the Rayleigh test of uniformity is noted below and written in bold if significant. ***G***, Mean dependent probabilities for the four possible outcomes of near-threshold trials given a time interval since the previous R-peak. The numbers for one time interval do not add up exactly to 100% across confident/unconfident misses and hits because of rounding and showing the mean across participants. The asterisks between the bars for confident hits indicate significant FDR-corrected *t*-tests.

Second, we repeated the analysis by splitting near-threshold trials based on the reported decision confidence. The unimodal distribution was also present for confident hits (*R* = 0.38, *p* = 0.002) but not for unconfident hits (*R* = 0.10, *p* = 0.69). Eighteen participants had a mean angle for confident hits in the last quarter of the cardiac cycle (270-360°). Confident misses also showed a unimodal distribution (*R* = 0.28, *p* = 0.04). Unconfident misses (*R* = 0.22, *p* = 0.14), confident correct rejections (*R* = 0.005 *p* = 1.00) and unconfident correct rejections (*R* = 0.01, *p* = 1.00) did not support rejecting the null hypotheses of a uniform distribution. Two participants were excluded from the analysis of unconfident correct rejections due to zero unconfident correct rejections (mean *n* = 20, *SD* = 22, range: 0-88).

Third, near-threshold trials were analyzed regarding their dependent probabilities of the four possible outcomes (unconfident misses, confident misses, unconfident hits, confident hits) given one of four time intervals after the R-peak (0 - 200 ms, 200 - 400 ms, 400 - 600 ms, 600 - 800 ms). FDR-corrected *t*-tests between time intervals for each outcome resulted in significant differences only for confident hits between the last interval (600 -800 ms) and the three other intervals (0 - 200 ms: *p* = 0.012; 200 - 400 ms: *p* = 0.0008; 400 - 600 ms: *p* = 0.0012). The significant comparison of linear mixed-effects models (LMMs) without and with the cardiac cycle information (*χ²* = 11.2, *p* = 0.004), indicated that the cardiac cycle explained variance of confidence decisions beyond their relationship with hit/miss responses.

Metacognitive efficiency was assessed across the cardiac cycle with response-specific M-ratios for each interval based on Bayesian hierarchical models (Fleming et al., 2017) for yes/no-responses (mean number of no-responses per cardiac interval = [88, 90, 84, 67], *SD* = [25, 24, 25, 29]; mean number of no-responses per cardiac interval = [50, 48, 49, 38], *SD* = [22, 22, 21, 17]). The means of the group-level M-ratio posterior distribution for no-responses were all below 1 across the cardiac cycle (Figure 4; mean group-level posterior distribution M-ratio(no) = [0.59, 0.56, 0.59, 0.58], *SD* = [0.75, 0.52, 0.45, 0.34]) whereas this was not the case for yes-responses: While the mean of the group-level M-ratio posterior distribution for yes-responses was below 1 at 0 - 200 ms after the R-peak, it was above 1 at 200 - 400 ms, and stabilized a bit below 1 at 400 - 800 ms (mean group-level posterior distribution M-ratio(yes) = [0.81, 1.05, 0.93, 0.95], *SD* = [0.36, 0.09, 0.23, 0.15]).

**Figure 4.**
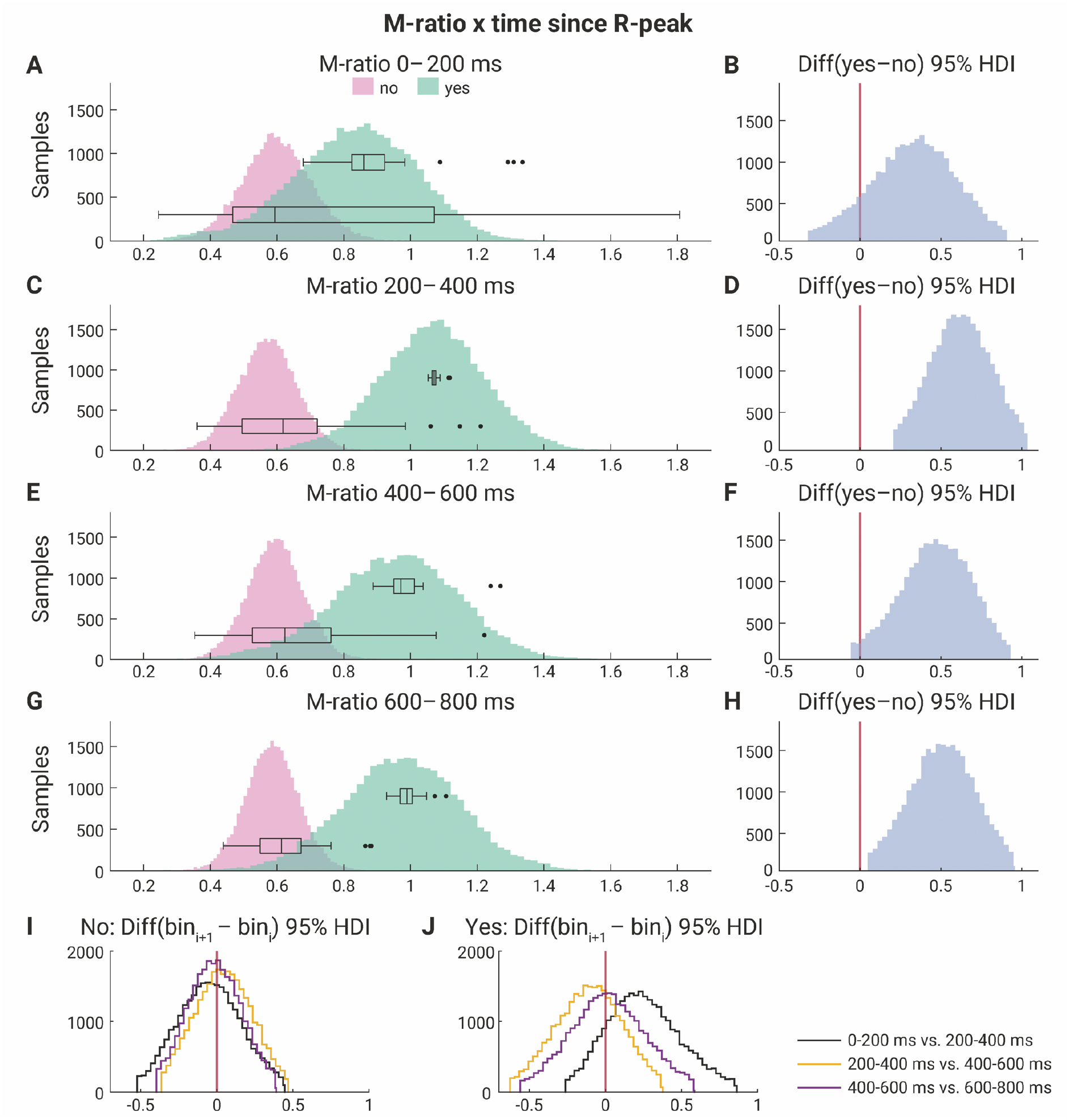
Response-specific metacognitive efficiency (M-ratios) across the cardiac cycle. At four cardiac intervals after the R-peak (***A***, ***C***, ***E***, ***G***) the posterior distributions of group-level M-ratios are shown for no (pink; correct rejection vs. miss) and yes-responses (green; hit vs. false alarm). On top of these histograms of Markov chain Monte Carlo (MCMC) samples, boxplots represent the participant-level M-ratios for yes/no-responses. M-ratios of 1 indicate that confidence ratings can perfectly discriminate between correct and incorrect responses. M-ratios below 1 indicate inefficient metacognition. The second column shows the difference between the posterior distributions of yes/no-responses as the 95% high-density intervals (HDI) at the four cardiac cycle intervals (***B***, ***D***, ***F***, ***H***). The last row shows the 95% HDIs between subsequent cardiac intervals (bin_i+1_ - bin*i*) for yes/no-responses (***I***, ***J***). These 95% HDIs indicate a credible difference between the corresponding group-level M-ratios if zero (red vertical line) is not included (***D***, ***H***).

For each cardiac cycle interval, the difference of the group-level M-ratio posterior distribution between yes/no-responses was assessed with 95% high-density intervals (HDI). For 200 - 400 ms and 600 - 800 ms, the 95% HDIs did not entail zero, thus providing evidence that metacognitive efficiency was higher for yes compared to no-responses. When comparing the group-level M-ratio posterior distribution between subsequent cardiac cycle intervals within yes/no-responses, the 95% HDIs always entailed zero. Hence, there was no statistical evidence for an overall modulation of metacognition across the cardiac cycle.

### Cardiac interbeat interval

For each stimulus-response condition (hit, miss, and correct rejection), we extracted the interbeat interval entailing the stimulus onset, as well as the two preceding and subsequent interbeat intervals (Figure 5A). We used a two-way repeated measures ANOVA to test the factors time and stimulus-response condition, and their interaction on interbeat intervals. The main effect of time (*F*(2.73,109.36) = 35.60, *p* = 3 x 10^-15^) and the interaction of time and stimulus-response condition (*F*(4.89,195.55) = 4.92, *p* = 0.0003) were significant. There was no significant main effect of stimulus-response condition on interbeat intervals (*F*(1.42,56.75) = 2.35, *p* = 0.12). Following, post-hoc *t*-tests were calculated at each interbeat interval between the stimulus-response conditions (5 x 3) and within each stimulus-response condition between subsequent interbeat intervals (3 x 4), resulting in 27 FDR-corrected *p*-values. At “S+1”, the interbeat intervals (IBI) for hits were significantly longer than for misses (*ΔIBI* = 5.2 ms, FDR-corrected *p* = 0.024) and correct rejections (*ΔIBI* = 4.4 ms, FDR-corrected *p* = 0.017). The interbeat intervals between misses and correct rejections did not differ significantly (FDR-corrected *p* = 0.62). At “S+2”, the interbeat intervals for hits were still longer compared to correct rejections (*ΔIBI* = 5.3 ms, FDR-corrected *p* = 0.014) but not to misses (FDR-corrected *p* = 0.25). Within each stimulus-response condition (hit, miss, and correct rejection) subsequent interbeat intervals differed significantly (FDR-corrected *p* < 0.005). Furthermore, the heart slowing ratio in all trials as well as in correct rejections (interbeat interval “Stimulus” to “S+1”) was correlated with near-threshold detection rate. This resulted in a strong correlation of heart slowing and detection task performance across participants (all trials: *r* = 0.53, *p* = 0.0004; correct rejections: *r* = 0.58, *p* = 0.00006).

**Figure 5.**
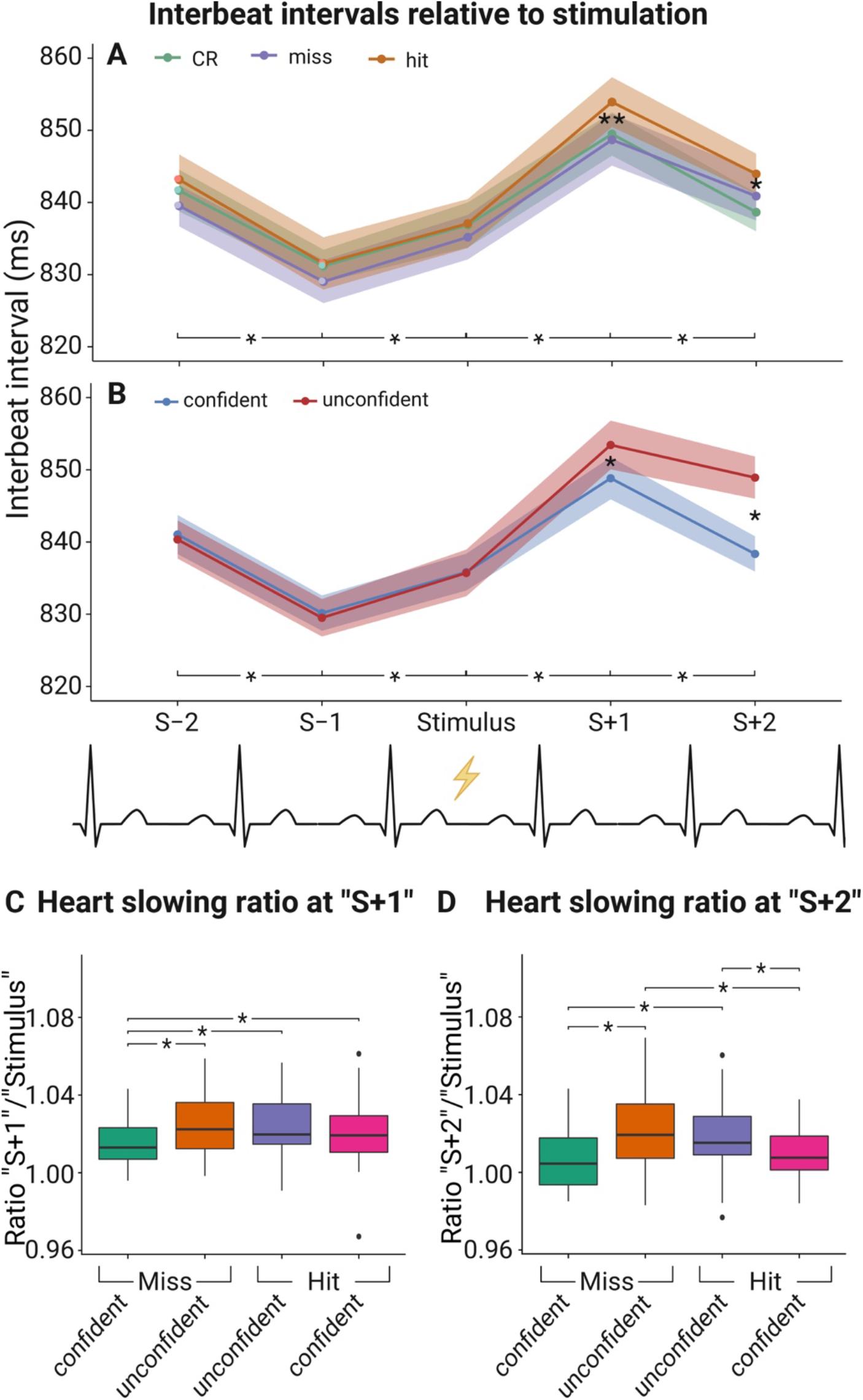
Interbeat intervals before and after the stimulus onset for (***A***) correct rejections (green), misses (purple), and hits (orange), and for (***B***) confident (blue) and unconfident (red) decisions. Confidence bands reflect within-participant 95% confidence intervals. The label “Stimulus” on the y-axis indicates the cardiac cycle when the stimulation or cue only were present. The labels “S-1” and “S+1” indicate the preceding and following intervals respectively. In ***A***, the two asterisks “**” at “S+1” indicate significant FDR-corrected *t*-tests between hits and misses, and between hits and correct rejections. The one asterisk “*” at “S+2” in ***A*** indicates a significant FDR-corrected *t*-test between hits and correct rejections. In ***B***, the asterisks at “S+1” and “S+2” indicate significant FDR-corrected *t*-tests between confident and unconfident decisions. The lines with asterisks on the bottom indicate significant FDR-corrected *t*-tests for subsequent interbeat intervals within all conditions. In ***C*** and ***D***, the ratio of interbeat intervals at “S+1” and “S+2” relative to “Stimulus” are shown for unconfident / confident misses and hits. The boxplots indicate the median (centered line), the 25%/75% percentiles (box), 1.5 times the interquartile range or the maximum value if smaller (whiskers), and outliers (dots beyond the whisker range). The asterisks between the boxplots indicate significant FDR-corrected *t*-tests.

To determine whether the longer interbeat intervals at “S+1” for hits as compared to misses and correct rejections, was due to longer systole or diastole, we automatically detected the T-wave end to separate both cardiac phases for statistical comparison. At “S+1”, the length of systoles was on average 324 ms (*SD* = 24 ms), and the length of diastoles 535 ms (*SD* = 107). Only diastoles for hits were significantly longer compared to misses (*ΔIBI* = 6.3 ms, FDR-corrected *p* = 0.006) and correct rejections at “S+1” (*ΔIBI* = 5.1 ms, FDR-corrected *p* = 0.006). Systoles at “S+1” showed no significant differences between the three conditions (hit vs. miss: FDR-corrected *p* = 0.81; hit vs. CR: FDR-corrected *p* = 0.81; miss vs. CR: FDR-corrected *p* = 0.81).

Furthermore, interbeat intervals of trials with confident and unconfident decisions independent of stimulus presence and yes/no-response (excluding false alarms) were compared with a two-way repeated measures ANOVA (Figure 5B). The main effects time (*F*(2.71, 108.31) = 42.37, *p* = 2 x 10^-16^) and confidence (*F*(1,40) = 5.36, *p* = 0.026), as well as the interaction of time and confidence were significant (*F*(2.73, 109.05) = 30.79, *p* = 2 x 10^-16^). Post-hoc *t*-tests between the two confidence categories for each interbeat interval (1 x 5) and within each confidence category between subsequent interbeat intervals (2 x 4) revealed significant longer post-stimulus interbeat intervals for unconfident compared to confident decisions at “S+1” (*ΔIBI* = 4.6 ms, FDR-corrected *p* = 0.005) and “S+2” (*ΔIBI* = 10.6 ms, FDR-corrected *p* = 6 x 10^-7^). All subsequent interbeat intervals differed significantly within each confidence category (FDR-corrected *p* < 0.05). When repeated for near-threshold trials only, the difference between confidence categories was still present within each awareness condition: unconfident hits and misses showed longer interbeat intervals at “S+2” compared to confident hits (*ΔIBI* = 5.7 ms, FDR-corrected *p* = 0.047) and confident misses respectively (*ΔIBI* = 10.2 ms, FDR-corrected *p* = 0.008).

Additionally, we tested whether the post-stimulus heart slowing ratio (interbeat interval “S+1” and “S+2” relative to “Stimulus”) differed in near-threshold trials between detection and confidence (Figure 5C, D). This approach has the advantage to account for the preceding interbeat interval “Stimulus” when comparing the interbeat interval differences at “S+1” and “S+2”. A two-way repeated measures ANOVA showed a main effect between confidence categories (“S+1”: *F*(1, 40) = 11.27, *p* = 0.02; “S+2”: *F*(1, 40) = 28.98, *p* = 0.000003) but not for detection (“S+1”: *F*(1, 40) = 4.04, *p* = 0.051; “S+2”: *F*(1, 40) = 0.13, *p* = 0.73). At “S+1”, post-hoc *t*-tests showed significant lower heart slowing ratios relative to “Stimulus” for confident misses compared to unconfident misses (FDR-corrected *p* = 0.001), unconfident hits (FDR-corrected *p* = 0.00002), and confident hits (FDR-correct *p* = 0.04). At “S+2”, heart slowing ratios relative to “Stimulus” were significantly lower for confident misses compared to unconfident misses (FDR-corrected *p* = 0.0001) and unconfident hits (FDR-corrected *p* = 0.0002), and for confident hits compared to unconfident hits (FDR-correct *p* = 0.001) and unconfident misses (FDR-correct *p* = 0.0004).

### Pulse wave relative to electric cardiac cycle

Next to the electric cardiac cycle, we assessed whether stimulus detection was dependent on the pulse wave cycle measured at the left middle finger. Pulse wave peaks were located in the cardiac cycle by calculating the time to the preceding R-peak: the pulse wave transit time (PWTT) and the PWTT relative to its current cardiac cycle in degree. The PWTT was on average 405 ms (*SD* = 24 ms, range: 354 - 439 ms). The pulse wave peak occurred on average in the middle of the cardiac cycle (mean angle *M*_PWTT_ = 178°, *R* = 0.91, *p* = 0) after the mean angle of confident misses (*M*_Confident miss_ = 96°) and before the mean angle of confident hits (*M*_Confident hit_ = 325°).

For putting the observed correlations between detection and the cardiac cycle in relation to the pulse wave peak, the analysis of near-threshold hit rates during different stimulus onset intervals after the R-peak was repeated limited to 0 - 400 ms with shorter intervals (50 ms) and without splitting by confidence. A one-way repeated-measures ANOVA showed a main effect by time interval on near-threshold detection rate (*F*(5.64, 225.46) = 3.15, *p* = 0.007). The near-threshold hit rates were significantly decreased before the pulse wave peak (mean PWTT = 405 ms) during the interval of 250 - 300 ms compared to the interval of 0 - 50 ms (FDR-corrected *p* = 0.038). The interval 250 - 300 ms was plotted on the average pulse wave locked to the preceding R-peak and its slope (difference between adjacent samples, first derivative) to determine the onset of the pulse wave arrival in the finger. The first derivatives of participant’s mean pulse waves showed that after 250 ms the pulse wave slopes substantially increased indicating the onset of the arriving pulse waves (Figure 6B).

**Figure 6.**
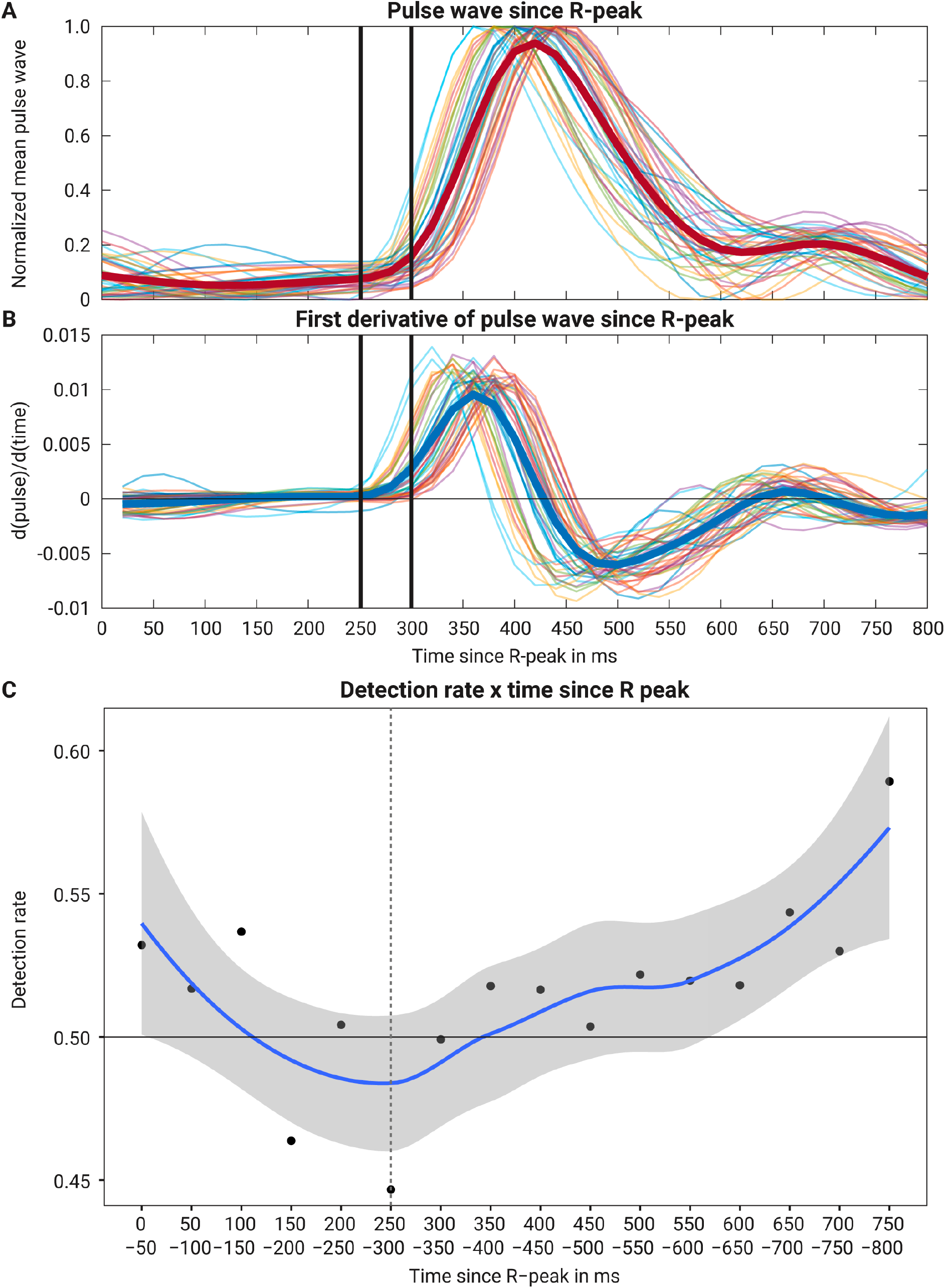
Pulse wave and detection relative to cardiac cycle. (***A***) Mean pulse waves measured at the left middle finger across all participants (red thick line) and for each participant (colored thin lines) locked to preceding R-peak. (***B***) First derivative of the mean pulse waves indicating the onset of the arriving pulse wave in the finger. The time window with the lowest detection rate is indicated with vertical thick black lines. (***C***) Detection rate of near-threshold trials in 50-ms stimulus onset intervals since preceding R-peak. The black dots indicate the mean across participants. The blue line is the locally smoothed loess curve with a 95% confidence interval (grey) across these means.

### Respiratory cycle

First, we investigated whether conscious tactile perception depends on the stimulus onset relative to the respiratory cycle. Thus, we calculated the mean angles for hits, misses, and correct rejections for each participant and tested their circular distribution with the Rayleigh test of uniformity. For all conditions, uniformity was rejected in favor of an alternative unimodal distribution (correct rejections: *R* = 0.58, *p* = 4 x 10^-7^; misses: *R* = 0.54, *p* = 2 x 10^-6^; hits: *R* = 0.65, *p* = 1 x 10^-8^; Figure 7). These unimodal distributions were centered at stimulus onset for the three conditions (mean angle *M*_correct rejection_= 3.2°, *M*_miss_ = 5.0°, and *M*_hit_ = 15.1°). Furthermore, we analyzed the circular distribution for each participant and stimulus-response condition. For hits, 38 of 41 participants showed a significant Rayleigh test after FDR-correction. For misses, 30 participants had a significant Rayleigh test, and for correct rejections, 32 participants. To assess whether the strength of the respiration locking differed significantly between hits, misses, and correction rejections, the circular variance of stimulus onset angles across trials was calculated for each stimulus-response condition and compared with *t*-tests. Hits had a lower circular variance than misses (*ΔV* = -0.044, *t*(40) = -3.17, *p* = 0.003) and correct rejections (*ΔV* = -0.035, *t*(40) = - 2.78, *p* = 0.008), i.e., exhibited a stronger clustering around the mean direction. There was no significant difference in circular variance between misses and correct rejections (*p* = 0.44). Furthermore, the circular variance of respiration phases in all trials as well as in correct rejections showed a negative medium correlation with near-threshold detection rate across participants (all trials: *r* = -0.38, *p* = 0.013; correct rejections: *r* = -0.41, *p* = 0.008).

**Figure 7.**
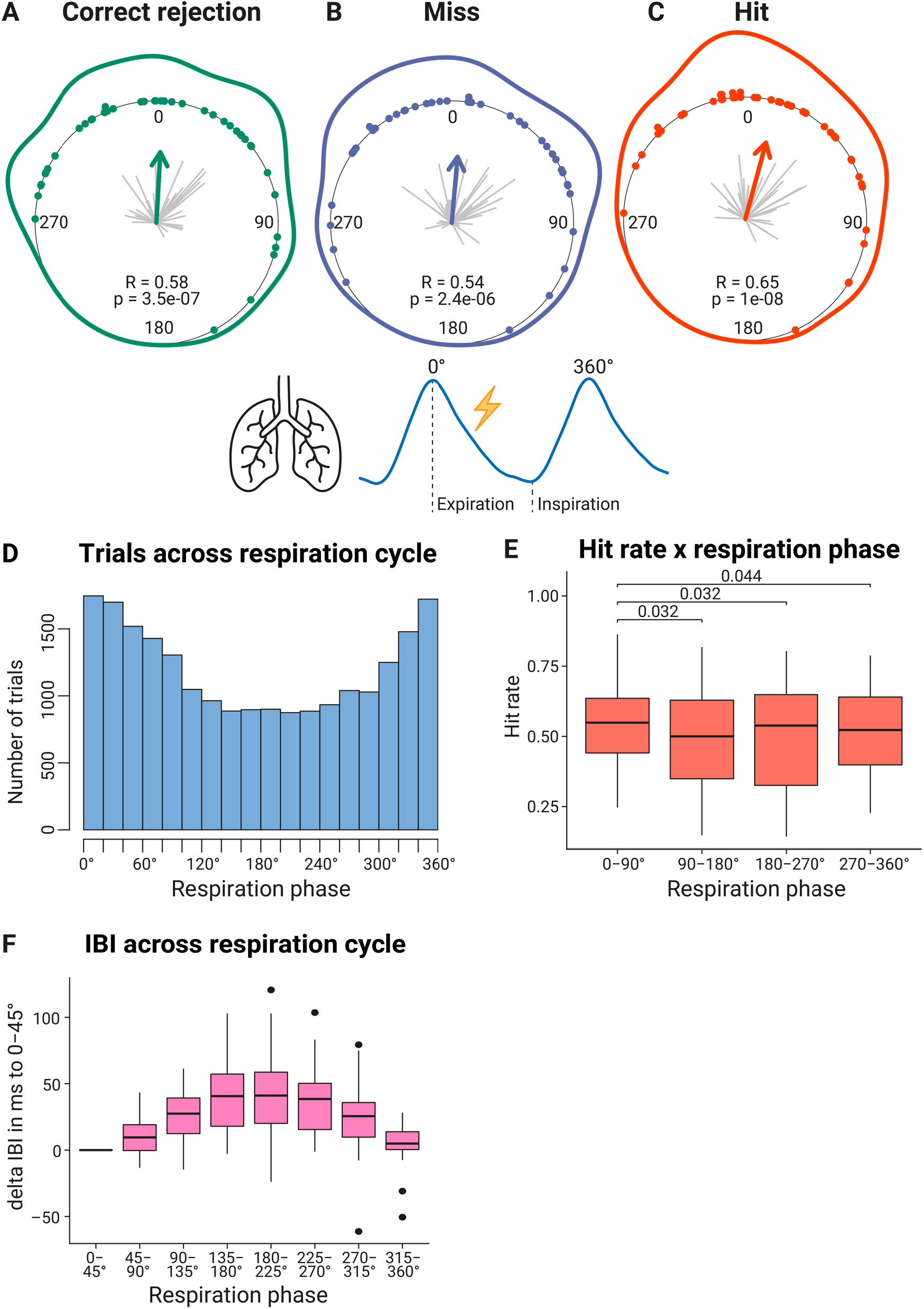
Circular distribution of mean stimulus onsets relative to the respiratory cycle for (***A***) correct rejections (green), (***B***) misses (purple), and (***C***) hits (red). Zero degree corresponds to expiration onset. Each dot indicates the mean angle of one participant. The grey lines originating in the center of the inner circle represent the resultant lengths *R_i_* for each participant’s mean angle. A longer line indicates a less dispersed intra-individual distribution (*V_i_* = 1-*R_i_*). The direction of the arrow in the center indicates the mean angle across the participants while the arrow length represents the mean resultant length *R*. The line around the inner circle shows the density distribution of these mean angles. The resulting p- value of the Rayleigh test of uniformity is noted below. ***D***, Histogram of respiration phases. Cumulative number of trials across all trials and participants for the relative position of the stimulus onset within the respiratory cycle binned in 20°-intervals from 0° to 360°. The Rayleigh test across all trials and participants was significant (*R* = 0.18, *p* = 2 x 10^-291^). ***E***, Detection rates for each quadrant of the respiratory cycle. Lines with *p*-values above the boxplots indicate significant FDR-corrected *t*-tests of all possible combinations. ***F***, Interbeat interval (IBI) differences for each eighth of the respiratory cycle relative to the first eighth (0-45°). The boxplots (***E***, ***F***) indicate the median (centered line), the 25%/75% percentiles (box), 1.5 times the interquartile range or the maximum value if smaller (whiskers), and outliers (dots beyond the whisker range).

We tested whether detection rates differed along the respiratory cycle. Thus, we binned near-threshold trials based on their relative position within the respiratory cycle in four quadrants (0-90°, 90-180°, 180-270°, and 270-360°), a one-way repeated-measures ANOVA was significant for the main effect quadrant on near- threshold detection rate (*F*(2.64,105.44) = 3.69, *p* = 0.018). Post-hoc *t*-tests revealed that only the first quadrant (0-90°) showed significantly greater hit rates (HR) compared to all other quadrants (90-180°: *ΔHR* = 3.8%, FDR-corrected *p* = 0.03; 180-270°: *ΔHR* = 3.7%, FDR-corrected *p* = 0.03; 270-360°: *ΔHR* = 2.3%, FDR-corrected *p* = 0.04).

Comparing cardiac interbeat intervals between eight 45°-intervals across the respiratory cycle showed a significant increase of interbeat intervals starting with the onset of expiration (increase from 0° to 225°), and a decrease starting with the onset of inspiration (decrease from 225° to 360°; Figure 7E). The inspiration onset was on average at 211° (range: 187-250°) within the respiratory cycle which started with the expiration onset (*R* = 0.97, *p* = 3 x 10^-16^).

Second, the distribution of mean angles was assessed for confident and unconfident decisions. Hits, misses, and correct rejections were split by decision confidence and the resulting distributions were evaluated with the Rayleigh test for uniformity. All stimulus-response conditions showed for unconfident and confident decisions a significant unimodal distribution locked around the stimulus onset: unconfident correct rejections (mean angle *M*_unconf_CR_ = 18.3°; *R* = 0.41, *p* = 0.001), confident correct rejections (*M*_conf_CR_ = 2.2°; *R* = 0.58, *p* = 4 x 10^-7^), unconfident misses (*M*_unconf_miss_ = 13.4°; *R* = 0.45, *p* = 0.0001), confident misses (*M*_conf_miss_ = 6.7°, *R* = 0.51; *p* = 0.00001), unconfident hits (*M*_unconf_hit_ = 10.4°; *R* = 0.51, *p* = 0.0001), and confident hits (*M*_conf_hit_ = 15.2°; *R* = 0.67, *p* = 4 x 10^-9^). Two participants had zero unconfident correct rejections and were not considered in the respective Rayleigh test.

Third, in order to examine aforementioned phase effects further, we investigated whether participants adjusted their respiration rhythm to the paradigm in the beginning of the experiment. Thus for the first 30 trials of the first block, a random-intercept linear regression model with maximum likelihood estimation (lmer function in R) was calculated to evaluate the effect of trial number on trial angle difference from the mean for each participant. The angle difference was determined between the stimulus onset angle within the respiratory cycle of each trial and the mean of all angles in the first block (“diff_angle2mean”). This analysis included 37 participants with a first block and excluded trials with false alarms. Comparing the model “diff_angle2mean ∼ 1 + trial + (1|participant)” with a random intercept-only model “diff_angle2mean ∼ 1 + (1|participant)” revealed an effect of trial on the difference to the angle mean within the first 30 trials of the first block (*χ²* = 5.84, *p* = 0.016). The fixed-effect slope was *b_1_* = -0.47 and the mean of the random-intercepts *b_0_* = 79.7 (*diff_angle2mean* = *b1* * *trial* + *b_0_*).

### Respiratory cycle duration

Given the previously reported heart slowing during conscious tactile perception (Motyka et al., 2019), we tested whether a similar effect was also present in the respiratory rhythm. Indeed, the mean duration of respiratory cycles differed between response categories (Figure 8), as indicated by a one-way repeated-measures ANOVA (*F*(1.49, 59.61) = 13.11, *p* = 0.0001). Post-hoc *t*-tests showed that respiratory cycles accompanying misses (mean *t* = 3.86 s) were significantly longer than respiratory cycles with correct rejections (mean *t* = 3.82 s, *Δt* = 40 ms, FDR-corrected *p* = 0.002) and with hits (mean *t* = 3.77 s, *Δt* = 91 ms, FDR-corrected *p* = 0.0002). Respiratory cycles with hits were also significantly shorter than correct rejections (*Δt* = 50 ms, FDR-corrected *p* = 0.014).

**Figure 8.**
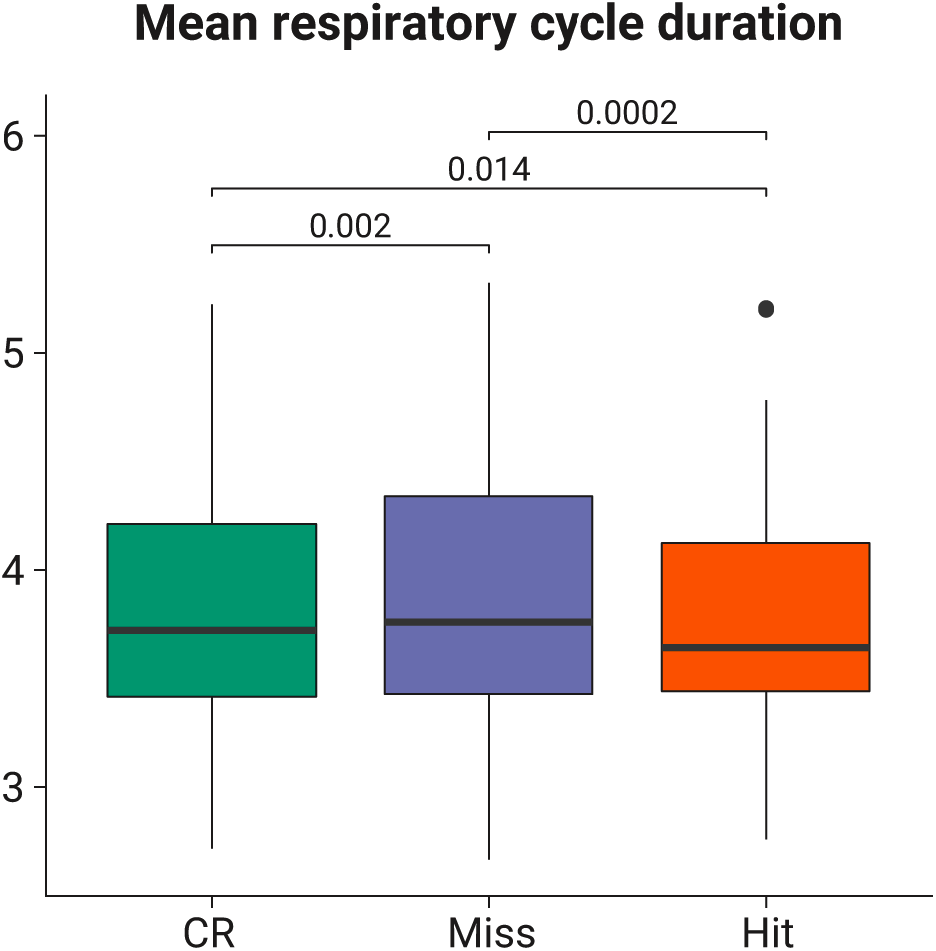
Mean respiratory cycle duration in seconds for correct rejections (green), misses (purple), and hits (red). The boxplots indicate the median (centered line), the 25%/75% percentiles (box), 1.5 times the interquartile range or the maximum value if smaller (whiskers), and outliers (dots beyond the whisker range). Significant post-hoc *t*-tests are indicated above the boxplot with a black bar and the respective FDR-corrected *p*-value.

Additionally, we analyzed whether the respiratory cycle duration differed between confident and unconfident hits and misses. There was a main effect by detection (*F*(1, 40) = 14.64, *p* = 0.0004) but not by confidence (*F*(1, 40) = 1.15, *p* = 0.29) on respiratory cycle duration in a two-way repeated measures ANOVA. The interaction of detection and confidence was not significant (*F*(1, 40) = 0.83, *p* = 0.37). Furthermore, we determined the expiration and inspiration duration for each respiratory cycle and compared them between hits, misses, and correct rejections. A two-way repeated measures ANOVA showed significant main effects of respiration phase (expiration longer than inspiration: *F*(1, 40) = 125.03, *p* = 7 x 10^-14^) and stimulus-response condition (*F*(1.45, 58) = 12.25, *p* = 0.00002). The interaction of respiration phase and stimulus-response condition was not significant (*F*(1.58, 63.15) = 0.22, *p* = 0.75). None of the six post-hoc *t*-tests between stimulus-response conditions for each respiration phase was significant after FDR-correction. The uncorrected *p*-values did not show evidence that the respiratory cycle duration differences were caused by the expiration or inspiration phase (expiration - correct rejection vs. miss: *p* = 0.04; correct rejection vs. hit: *p* = 0.21; miss vs. hit: *p* = 0.02; inspiration - correct rejection vs. miss: *p* = 0.09; correct rejection vs. hit: *p* = 0.06; miss vs. hit: *p* = 0.01).

### Phase-locking between cardiac and respiratory activity

Due to the natural coupling of cardiac and respiratory rhythms (Dick et al., 2014), we investigated whether phase-locking of both rhythms is associated with conscious tactile perception. Phase-locking values (PLVs) were calculated across trials using n:m synchronization (Tass et al., 1998; Lachaux et al., 1999) to account for the different frequency bands of the two signals. PLVs were compared between hits, misses, and correct rejections with a one-way repeated-measures ANOVA. The ANOVA showed no significant main effect of stimulus-response condition on PLVs between cardiac and respiratory activity (*F*(1.98,79.15) = 1.72, *p* = 0.19).

### Peripheral nerve activity

For the sub-sample of twelve participants with peripheral nerve recordings at the left upper arm, there was no somatosensory evoked potential associated with near-threshold stimuli. Also, the grand mean across participants did not show a difference between trials with and without near-threshold stimulation. We concluded that near-threshold stimulation intensities (in the given sub-sample on average 1.88 mA, range: 0.79-2.50 mA) did not produce sufficiently high peripheral somatosensory evoked potentials to measure them non-invasively from the inner side of the upper arm. Hence, we did not further pursue the analysis of peripheral somatosensory evoked potentials. (Yet note that peripheral somatosensory evoked potentials were observed in a pilot study with the same acquisition setup but applying super-threshold stimulation intensities of 6 mA.)

## Discussion

In this study, we confirm our previous finding that stimulus detection varies along the cardiac cycle (Motyka et al., 2019; Al et al., 2020, 2021). With the additional recording of photoplethysmography, decision confidence, and respiratory activity, we obtain several new findings regarding the integration of cardiac and respiratory signals in perceptual decision making: We precisely pinpoint the period of lowest tactile detection rate at 250 - 300 ms after the R-peak, and we show a variation of confidence ratings across the cardiac cycle. A further new finding is that confidence ratings are the major determinant of cardiac deceleration. We confirm previous findings of an alignment of the respiratory cycle to the task cycle and we observed that this alignment follows closely the modulation of heart frequency (HF) across the respiratory cycle (sinus arrhythmia) with preferred stimulation onsets during periods of highest HF. Detection rate was highest in the first quarter of the respiratory cycle (after expiration onset), and temporal clustering during the respiratory cycle was more pronounced for hits than for misses and - interindividually - stronger respiratory phase-locking was associated with higher detection rates. Taken together, our findings show how tuning to respiration and closely linked cardio-respiratory signals are integrated to achieve optimal task performance.

### Detection varies across the cardiac cycle and is lowest 250 - 300 ms post R-peak

While replicating the unimodal distribution of hits within the cardiac cycle here for the third time in an independent study of somatosensory detection (Motyka et al., 2019; Al et al., 2020, 2021), we now located the decreased near-threshold detection rate more precisely 250 - 300 ms after the R-peak, before the pulse wave peak (405 ms) in the middle of the cardiac cycle (178°). The slope of the pulse wave showed a take-off around 250 ms after the preceding R-peak, indicating the onset of the pulse wave arrival. The explanation of the cardiac cycle effect on somatosensory detection stays speculative. In our previous study (Al et al., 2020), we found this systolic suppression to be associated with a change in sensitivity and with a reduction of the P300 SEP component which is commonly assumed to encode prediction (errors) (Friston, 2005). We therefore postulated that the prediction of the pulse-wave associated peripheral nerve activation (Macefield, 2003) also affects the perception of other weak stimuli in the same time window since they are wrongly assumed to be a pulse-related ‘artifact’. Our new finding of temporally locating the lower tactile detection at the pulse wave onset and not during maximal peripheral vascular changes in the finger further supports this view. Perception of heartbeats has been reported to occur in the very same time interval of 200 - 300 ms after the R-peak (Yates et al., 1985; Brener and Kluvitse, 1988; Ring and Brener, 1992). While this temporal judgement is unlikely to be solely based on the pulse wave in the finger - heartbeat sensations were mainly localized on the chest (Khalsa et al., 2009; Hassanpour et al., 2016) - it is consistent with the prediction of strongest heartbeat-related changes at 250 - 300 ms and an attenuated detection of weak stimuli presented in the same time window.

### Confidence ratings vary across the cardiac cycle

We also show that the cardiac cycle had a relationship with confidence ratings, in addition to the association of hit/miss responses with confidence ratings. When comparing the dependent probabilities of the four possible outcomes in near-threshold trials (unconfident/confident miss/hit) – only the number of confident hits increased at the end of the cardiac cycle (600 - 800 ms) compared to 0 - 600 ms.

By determining M-ratios relating meta-*d’* to *d’*, we show that metacognitive efficiency for yes-responses is generally higher than for no-responses (close to 1 (“optimal”) versus below 1 (“inefficient”)). In our data, there was no evidence for an overall modulation of metacognition across the cardiac cycle. Qualitatively, while metacognition for no-responses is clearly smaller than 1 (“inefficient”) for all cardiac intervals, metacognition for yes-responses seems to be shifted below 1 (“inefficient”) at the onset of systole (0 - 200 ms after R-peak), shifted above 1 (“super-optimal”) during the period of 200 - 400 ms, and stabilizing over 1 at 400 - 800 ms. Future research must show whether this qualitative observation can be replicated. If so, the systolic variation might be related to a decisional conflict during an interval with the highest uncertainty whether a weak pulse was generated internally (heartbeat) or applied externally (Allen et al., 2019).

The higher confidence ratings for misses than for hits are most likely due to the higher expectation of no-responses which is about 66% – given 1/3 null trials and 2/3 ‘50% near-threshold’ trials. For visual decision-making, confidence has been shown to be influenced by probabilities and - hence - expectations that a stimulus would occur (Sherman et al., 2015) and that the decision would be correct (Aitchison et al., 2015). In an independent fMRI study with near-threshold somatosensory stimuli using a four-point confidence scale, we equally found lower confidence for hits than for misses (Grund et al., 2021). It is not as straightforward to explain the overall lower metacognitive efficiency for no-responses versus yes-responses. We speculate that it is probably related to the different likelihoods of the respective decisional alternatives: Among the no-responses ‘Miss’ and ‘Correct rejection’ have an almost equal likelihood while among yes-responses ‘Hits’ are much more likely than ‘False Alarms’.

### Heart slowing and perceptual decision making

In the present study, we confirmed cardiac deceleration related to the parasympathetic correlate of the orienting response to a change in the environment (Sokolov, 1963) for all trial types even for trials without stimulation and we also confirmed the previously reported more pronounced heart deceleration with conscious perception was replicated (Park et al., 2014; Cobos et al., 2019; Motyka et al., 2019). Interestingly, however, when confidence ratings were taken into account this effect was reduced (particularly at two interbeat intervals after the stimulation) and our findings indicate that heart rate slowing is mainly due to confidence rating, such that unconfident decision are associated with stronger heart rate slowing. Increased heart slowing for unconfident decisions might be associated with uncertainty, because heart slowing has been reported for the violation of performance-based expectations in a learning paradigm (Crone et al., 2003), for errors in a visual discrimination task (Łukowska et al., 2018), and for error keystrokes by pianists (Bury et al., 2019). Previous studies have linked heart rate changes with confidence. For example, in a visual discrimination task, confidence has been associated with heart acceleration which - in turn - attenuated the heart slowing caused by the orienting response to the stimulus (Allen et al., 2016). Since this effect was reversed by a subliminal and arousing negative emotional cue, confidence was interpreted as an integration of exteroceptive and interoceptive signals (Allen et al., 2016, 2019). A related phenomenon may underlie our results, in that rapid heart rate changes are transmitted upstream to be integrated in the decision process and particularly its metacognitive aspects.

Variations in the cardiac cycle were mainly due to changes in the length of diastoles which is in line with previous literature showing that cardiac cycle length is mainly modulated by diastole (Levick, 1991).

Interestingly, the extent of cardiac deceleration (we tested for all trials and for correct rejections) showed a positive correlation with near-threshold detection across participants. This is the case despite the fact that in all participants near-threshold stimulation intensity was adjusted before the experiment such that they detected about 50% of the stimulus trials. We thus have to assume that from the starting point of about 50% detection rate (without adjustment of the respiration), those participants who had a more pronounced heart slowing during the experiment improved their detection rate more than other participants.

### Respiration locking and perceptual decision making

Localizing stimulus onsets in the respiratory cycle revealed that (expected) stimulus onsets were locked to respiration. During the first thirty trials, the angular difference of onset time points to the mean angle showed a linear decrease as participants adapted their respiration rhythm to the paradigm. Intra-individual circular variance of stimulation onsets was lower for hits than misses, indicating a more pronounced respiratory phase-clustering went along with a higher likelihood of hits. Hit rates were greater in the first quadrant after expiration onset compared to all other three quadrants of the respiratory cycle. Furthermore, there was a negative correlation between intra-individual circular variance of respiration phases in all trials (and also correct rejections) and near-threshold detection rate across all participants. Together these results suggest that respiration-locking is beneficial for task performance.

Interestingly, the frequency distribution of stimulation onsets closely matched heart rate changes along the respiratory cycle. Heart rate showed the well-known increase with expiration and decrease with inspiration (sinus arrhythmia), and stimulus onsets occurred most frequently during the period with shortest interbeat intervals i.e., highest heart rate. It is known that not only heart rate but also neural excitability changes during the respiratory cycle. In a recent study, the course of alpha power – known to be inversely related to excitability – across the respiratory cycle has been shown to have a minimum around expiration onset (Kluger et al., 2021) – the time period when in our study most stimuli were timed. Also for tactile stimuli, it has been shown that alpha power in central brain areas is related to conscious detection (Schubert et al., 2009; Nierhaus et al., 2015; Craddock et al., 2017; Forschack et al., 2020; Stephani et al., 2021). Taken together, respiration phase locking might be used to increase the likelihood to detect faint stimuli in a phase of highest cortical excitability (attention).

While the mean angle across all participants locked at expiration onset, participant’s individual mean angles ranged from late inspiration to early expiration (circa >270° and <90°). The inspiration onset was on average at 211°. Thus, the current data does not allow to determine whether participants tuned their inspiration or expiration onset. Possibly participants adapted their respiration rhythm to the cue onset which occurred 500 milliseconds before the stimulus onset. Given a mean respiratory cycle duration of 3.82 seconds, the cue onset was 47 degrees before the stimulus onset.

## Conclusion

The two predominant body rhythms modulate conscious tactile perception. Our data indicate that phase-locking of respiration facilitates perception by optimal timing of stimuli in periods of highest heart rate and cortical excitability. Tactile detection and related decision confidence also vary characteristically during the cardiac cycle and the effects seem best explained by an interoceptive predictive coding account which is meant to model and suppress bodily changes related to the heartbeat.

## Acknowledgements

The research was funded by the Max Planck Society. We thank Sylvia Stash for her data acquisition support; Mina Jamshidi Idaji and Carina Forster for data analysis advice and support; and Heike Schmidt-Duderstedt and Kerstin Flake for preparing the figures for publication.

## Notes

https://github.com/grundm/respirationCA

### Competing Interest Statement

The authors have declared no competing interest.

### Summary of Updates

Updated statistical analysis of metacognition

